# TGF-β2-induced Snail-mediated EndMT signaling relies on both Smad-dependent and independent pathways

**DOI:** 10.64898/2026.08.17.745236

**Authors:** Valerie S. Kalluri, Bingrui Li, Anais M. Comptdaer, Michelle Kirtley, Kent A. Arian, Xunian Zhou, Raghu Kalluri

## Abstract

Endothelial-to-mesenchymal transition (EndMT) has become a central mechanism in developmental biology, fibrosis, vascular disease, and cancer. We previously reported on an integrated signaling model in which TGF-β2 induces EndMT through coordinated activation of Smad-dependent and Smad-independent signaling pathways converging on Snail, while GSK-3β regulates Snail activity. We performed a systematic figure-by-figure reproducibility analysis of the original publication. Independent studies published between 2011 and 2026 were identified and curated according to predefined inclusion criteria. Each original experimental conclusion was evaluated for independent confirmation. In parallel, selected biochemical experiments were independently reproduced using newly acquired reagents and contemporary Western blot methodologies. Independent publications consistently reproduced each major mechanistic conclusion of the original study, including activation of Smad, ERK, PI3K/AKT, and p38 MAPK signaling, regulation of Snail expression, EndMT-associated marker switching, and GSK-3β-dependent control of Snail activity. Independent laboratory experiments reproduced the principal biochemical findings using contemporary reagents and experimental workflows. The combined literature analysis and independent laboratory replication demonstrate that the mechanistic framework in our previous study has remained reproducible across multiple laboratories, endothelial cell types, disease models, and fifteen years of investigation. This work illustrates a complementary framework for assessing reproducibility that integrates direct experimental replication with cumulative independent validation.

## Introduction

Reproducibility is widely recognized as a cornerstone of scientific progress. Over the past decade, concerns regarding the reproducibility of biomedical research have stimulated numerous initiatives aimed at independently validating influential findings. These efforts have generally focused on repeating experiments under controlled conditions within independent laboratories. These approaches provide valuable information regarding technical reproducibility, but they could underestimate the robustness of discoveries that have subsequently been reproduced independently using diverse experimental models and biological systems. Cumulative independent validation across time represents an additional dimension of reproducibility that has received comparatively little systematic study. Mechanistic discoveries frequently become integrated into the broader scientific literature through gradual independent confirmation rather than formal replication studies. Over time, investigators extend original observations into new experimental settings, employ distinct methodologies, and evaluate similar biological questions in different disease contexts. Collectively, these independent studies provide an important, although often underappreciated, measure of reproducibility. Assessing the cumulative reproducibility of influential mechanistic studies therefore represents a complementary approach to evaluating scientific rigor.

Endothelial-to-mesenchymal transition (EndMT) is now recognized as a fundamental mechanism regulating cardiovascular development, tissue fibrosis, vascular remodeling, and tumor progression^1-9^. During EndMT, endothelial cells lose characteristic endothelial features, including expression of VE-cadherin and CD31, while acquiring mesenchymal properties such as fibroblast-specific protein-1 (FSP-1), α-smooth muscle actin (α-SMA), and enhanced migratory capacity. EndMT contributes to numerous pathological processes, including cardiac fibrosis, pulmonary hypertension, diabetic nephropathy, atherosclerosis, ocular disease, and cancer-associated fibroblast formation^1-9^. We previously reported that TGF-β2 induces EndMT through coordinated activation of canonical Smad signaling together with MEK/ERK, PI3K/AKT, and p38 MAPK pathways^10^. The study further demonstrated that these signaling pathways converge upon regulation of the transcription factor Snail and established that inhibition of glycogen synthase kinase-3β (GSK-3β) is required for Snail to fully induce EndMT. These observations provided one of the earliest integrated mechanistic models explaining how multiple TGF-β signaling pathways cooperate to regulate endothelial plasticity. Since publication, EndMT has become an extensively investigated biological process, providing an opportunity to evaluate the long-term reproducibility of the original mechanistic model. Rather than assessing a single experimental replication, we sought to determine the extent to which each principal conclusion of the original study has been independently reproduced across the scientific literature. We further performed an independent laboratory replication of selected key biochemical experiments using newly acquired reagents and contemporary experimental workflows. Together, these complementary analyses provide a comprehensive assessment of the reproducibility of TGF-β2-induced EndMT signaling over fifteen years of subsequent investigation. Here, we systematically evaluated the reproducibility of the mechanistic conclusions we previously reported^10^ through two complementary approaches: (i) a structured figure-by-figure literature review documenting independent replication of each principal experimental finding, and (ii) direct laboratory reproduction of selected biochemical experiments using independently sourced reagents and current experimental standards.

## Materials and Methods

Experimental conditions, treatment durations, inhibitor concentrations, and analytical methods were matched to those described previously while utilizing currently available commercial reagents.

### Cell culture

Primary human cutaneous microvascular endothelial cells (HCMECs) were procured from Cell Systems (ACBRI 538, Passage 3, lot 538 01 01 01 2M) and cultured using the manufacturer’s culture reagents before transitioning to previously used endothelial growth conditions (EBM-2 culture media fully supplemented according to the manufacturer’s direction, Lonza CC-3156 lot 0001424029 and Lonza CC-4176 batch 00001449162). The cells were grown to reach approximately 80–90% confluence on dishes coated with 0.1% gelatin (Cell Biologics, Cat. 6950, Lot 0323). Twenty-four hours before experimental treatment, cells were transferred to serum-free endothelial medium (Gibco, 11111-044, lot 3221721). For pathway activation studies, recombinant human TGF-β2 was added at a final concentration of 10 ng/mL. Recombinant human TGF-β2 (R&D Systems, Cat. 302-B2-002/CF, Lot KF2125121) was used for all stimulation experiments. Cells were harvested after 30 minutes for phosphorylation studies or after 48 hours for morphological assessment (Keyence BZ-X1000) and EndMT marker analysis, consistent with the original experimental design. To assess the contribution of Smad-independent signaling pathways during TGF-β2-induced EndMT, cells were pretreated for 1 hour before TGF-β2 stimulation with SB202190 (p38 MAPK inhibitor; MedChemExpress, Cat. HY-10295, Batch 835589) at 25 μM, LY294002 (PI3K inhibitor; MedChemExpress, Cat. HY-10108, Batch 267411) at 50 μM, U0126 (MEK1/2 inhibitor; MedChemExpress, Cat. HY-12031A, Batch 840583) at 10 μM. For experiments evaluating GSK-3β inhibition, LiCl (Millipore Sigma, 310468, Batch MKCZ0556) was added at a final concentration of 20 mM at 24 hours following plasmid transfection, as described previously^10^.

### Suppression of Snail expression using siRNA

For Snail knockdown experiments, endothelial cells were transfected with Snail-specific siRNA or a non-targeting control siRNA using Lipofectamine 3000 transfection kit (Invitrogen, Cat. L3000015, Lot 2905459) according to the manufacturer’s instructions. Cells were subsequently treated with recombinant TGF-β2 for 48 hours before protein analysis. The experimental design reproduced the conditions used in the original publication to evaluate the requirement of Snail during EndMT. siRNA were purchased from Sigma-Aldrich. The following sequences were used: Snail sense: CCACAGAAAUGGCCAUGGGAAGGCCAC[dT][dT], Snail anti-sense: GUGGCCUUCCCAUGGCCAUUUCUGUGG[dT][dT], Scrbl sense: UCACAAGGGAGAGAAAGAGAGGAAGGA[dT][dT], Scrbl anti-sense: UCCUUCCUCUCUUUUCUCUCCUUgUGA[dT][dT]

### Plasmid transfection

For Snail overexpression studies, endothelial cells were transiently transfected with either pcDNA3 vector (Addgene, plasmid #74165) or pcDNA3-Snail (Addgene, plasmid #31697) expression plasmids. Twenty-four hours following transfection, cultures were treated with LiCl (20 mM) to inhibit GSK-3β signaling before collection for immunoblot analysis. For experiment using dominant negative Smad4 transfection, pCMV5 DPC4 (1-514) was used (Addgene, plasmid #14040). Transfection were carried out using Lipofectamine 3000 transfection kit (Invitrogen, Cat. L3000015, Lot 2905459) according to the manufacturer’s instructions.

### Western blot analysis

Cells were lysed using RIPA buffer (Pierce, Cat. 89900) supplemented with protease and phosphatase inhibitors (Roche, Cat. 4693116001). Lysates were cleared by centrifugation at 12,000*g* for 10min at 4◦C and heated for 5min at 95◦C with LDS sample buffer (ThermoFisher, Cat. NP0007). Bolt™ mini gels were used for electrophoresis (ThermoFisher) and protein were transferred to PVDF membranes using Trans-Blot Transfer System (Bio-Rad). The membranes were blocked for 40min to 1h in 5% milk in TBS-tween at RT. Primary antibodies were incubated overnight. Secondary antibodies (Horsradish peroxidase conjugated) were incubated for 40 min to 1h at RT. All antibodies were diluted in 3% BSA in TBS-tween.

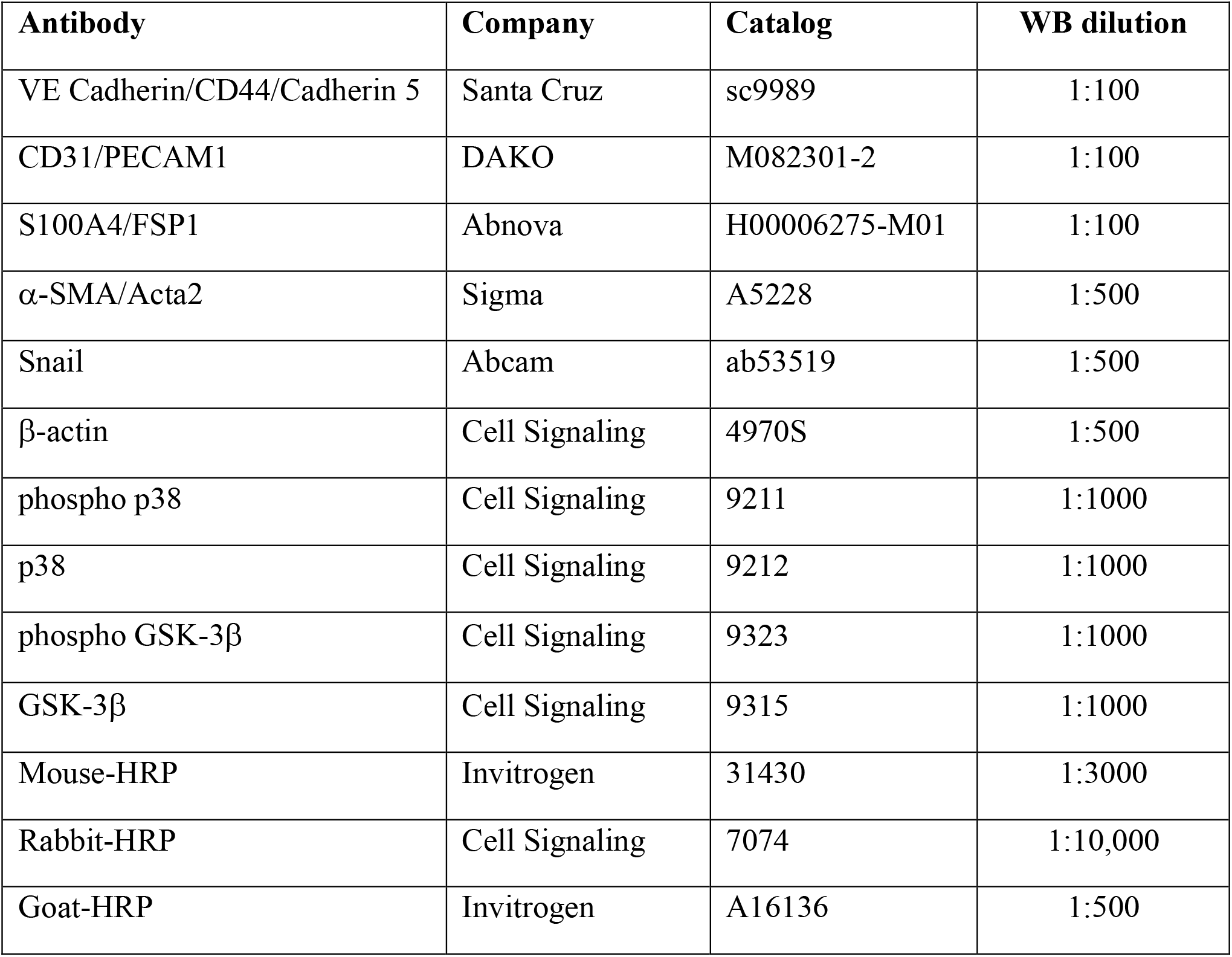

| **Antibody** | **Company** | **Catalog** | **WB dilution** |
| --- | --- | --- | --- |
| VE Cadherin/CD44/Cadherin 5 | Santa Cruz | sc9989 | 1:100 |
| CD31/PECAM1 | DAKO | M082301-2 | 1:100 |
| S100A4/FSP1 | Abnova | H00006275-M01 | 1:100 |
| α-SMA/Acta2 | Sigma | A5228 | 1:500 |
| Snail | Abcam | ab53519 | 1:500 |
| β-actin | Cell Signaling | 4970S | 1:500 |
| phospho p38 | Cell Signaling | 9211 | 1:1000 |
| p38 | Cell Signaling | 9212 | 1:1000 |
| phospho GSK-3β | Cell Signaling | 9323 | 1:1000 |
| GSK-3β | Cell Signaling | 9315 | 1:1000 |
| Mouse-HRP | Invitrogen | 31430 | 1:3000 |
| Rabbit-HRP | Cell Signaling | 7074 | 1:10,000 |
| Goat-HRP | Invitrogen | A16136 | 1:500 |

## Results

### Independent literature validates activation of multiple signaling pathways downstream of TGF-β2

Our previous study demonstrated that TGF-β2 activates canonical Smad signaling together with ERK1/2, PI3K/AKT, and p38 MAPK pathways in endothelial cells, and that pharmacologic inhibition of these pathways prevents EndMT^10^. To evaluate reproducibility of these findings, we performed a systematic figure-by-figure literature analysis encompassing publications from 2011 through 2026. For each experimental panel, independent studies were identified that directly reproduced either the biochemical observations or the mechanistic conclusions using independent investigators, endothelial cell models, and disease contexts. Multiple independent studies subsequently demonstrated activation of Smad signaling downstream of TGF-β2 during EndMT and confirmed the requirement for Smad-dependent regulation of Snail expression^11,12^. Likewise, independent investigations consistently demonstrated activation of ERK1/2 Ref^13,14^, PI3K/AKT^15-17^, and p38 MAPK^12,18-20^ signaling following TGF-β2 stimulation in endothelial cells, while pharmacological inhibition of these pathways suppressed EndMT-associated phenotypic changes^12,16,21-23^. Collectively, these studies reproduced the central signaling architecture originally proposed in 2011 across multiple endothelial cell types, including human microvascular, cardiac, corneal, and Schlemm’s canal endothelial cells.

### Independent laboratories confirm regulation of Snail by convergent TGF-β signaling pathways

The original publication demonstrated that Smad, ERK, PI3K, and p38 signaling cooperate to induce Snail expression and that inhibition of any individual pathway attenuates Snail induction. Subsequent investigations independently confirmed that TGF-β2-induced Snail expression requires coordinated signaling through canonical and non-canonical pathways^11-13,16,22^. Collectively these studies further established that Snail functions as a central downstream transcriptional regulator governing endothelial plasticity during EndMT. Although individual studies expanded upon the molecular mechanisms controlling Snail transcription and stability, the overall signaling framework remained highly consistent with the original mechanistic model.

### Independent studies reproduce Snail-dependent regulation of EndMT

We demonstrated that siRNA-mediated knockdown of Snail prevents TGF-β2-induced EndMT, whereas expression of Snail alone is insufficient to induce endothelial transition in the absence of appropriate post-translational regulation^10^. Numerous independent investigations subsequently confirmed that Snail is required for EndMT in diverse biological contexts including cardiac fibrosis, vascular remodeling, hypoxia-induced EndMT, and inflammatory signaling^22,24-27^. Suppression of Snail prevents EndMT marker switching^12,21,22,27-29^. Several studies additionally supported the concept that Snail protein activity is tightly regulated through post-translational mechanisms, explaining why Snail expression alone may be insufficient to induce complete EndMT in specific endothelial cell populations^30-32^. These observations collectively reinforce the concept that both transcriptional induction and post-translational regulation of Snail are necessary components of endothelial phenotypic conversion.

### Independent experimental replication confirms TGF-β2 promotes Snail-mediated EndMT through convergence of Smad-dependent and Smad-independent signaling

To complement the literature analysis, we independently repeated selected biochemical experiments from the original publication using newly acquired antibodies, independently sourced reagents, and contemporary Western blot methodologies. TGF-β2-induced phosphorylation of p38 MAPK and this is inhibited by the p38α and p38β isoforms of mitogen-activated protein (MAP) kinases SB202190 (**Figure 1, Supplementary Fig. 1A**), reproducing the biochemical findings previously reported (Figure 1D in Ref^10^). The regulation of endothelial (VE-cadherin, CD31) and mesenchymal (FSP-1, α-SMA) marker expression demonstrating TGF-β2 induced EndMT was specifically inhibited following Smad-dependent and Smad-independent signaling pathways suppression (**Figure 2, Supplementary Fig. 1B**), reproducing the principal observations from findings previously reported (Figure 2C in Ref^10^). Specifically, dominant-negative Smad4 (DN-Smad4), MEK1/2 inhibitor (U0126), PI3K inhibitor (LY294002), or p38 MAPK (SB202190) inhibitor prevented TGF-β2-induced EndMT morphological changes (**Figure 2A**) and EndMT marker expression changes (**Figure 2B, Supplementary Fig. 1B**). Likewise, siRNA-mediated inhibition of Snail prevented EndMT-associated marker switching (**Figure 3, Supplementary Fig. 2**), reproducing our previously reported findings (Figure 3B in Ref^10^). Finally we previously reported that over expression of Snail (pcDNA3-Snail expression plasmid), in the context of inhibition of GSK-3β (using LiCl^33^), allowed Snail to induce EndMT (Figure 5 in Ref^10^). Snail over expression alone did not lead to GSK-3β phosphorylation, but Snail levels were increased with concomitant treatment with LiCl (**Figure 4A, Supplementary Fig. 3A**). Combined Snail overexpression and GSK-3β inhibition reproduced the original findings demonstrating that GSK-3β functions as a critical regulator of Snail activity during EndMT (**Figure 4B, Supplementary Fig. 3B**). Taken together, these results support that inhibition of GSK-3β allows for Snail mediated induction of EndMT.

**Figure 1.**
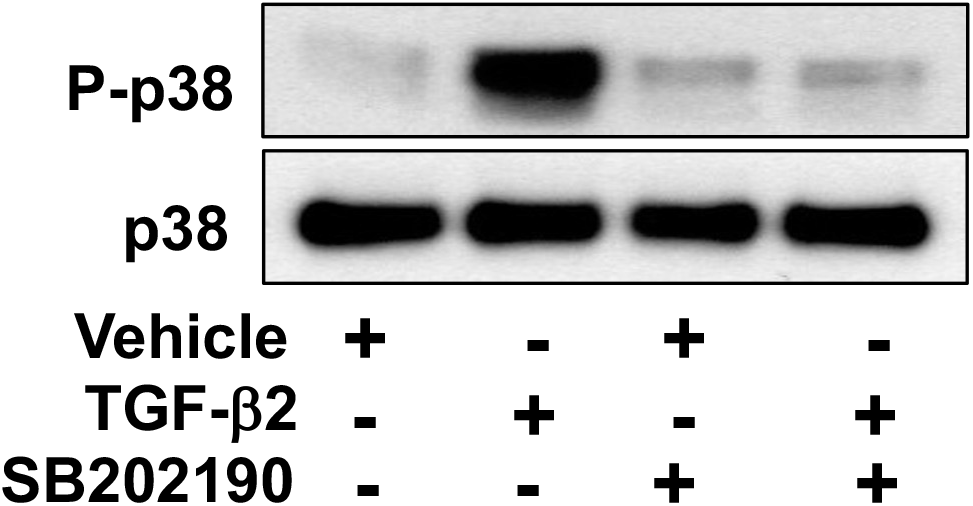
TGF-β2 activates p38 MAPK signaling in endothelial cells. Immunoblotting for phosphorylation levels of p38 MAPK and total p38 in endothelial cells under the listed conditions. TGF-β2: 10ng/mL; p38 inhibitor (SB202190), 25μM.

**Figure 2.**
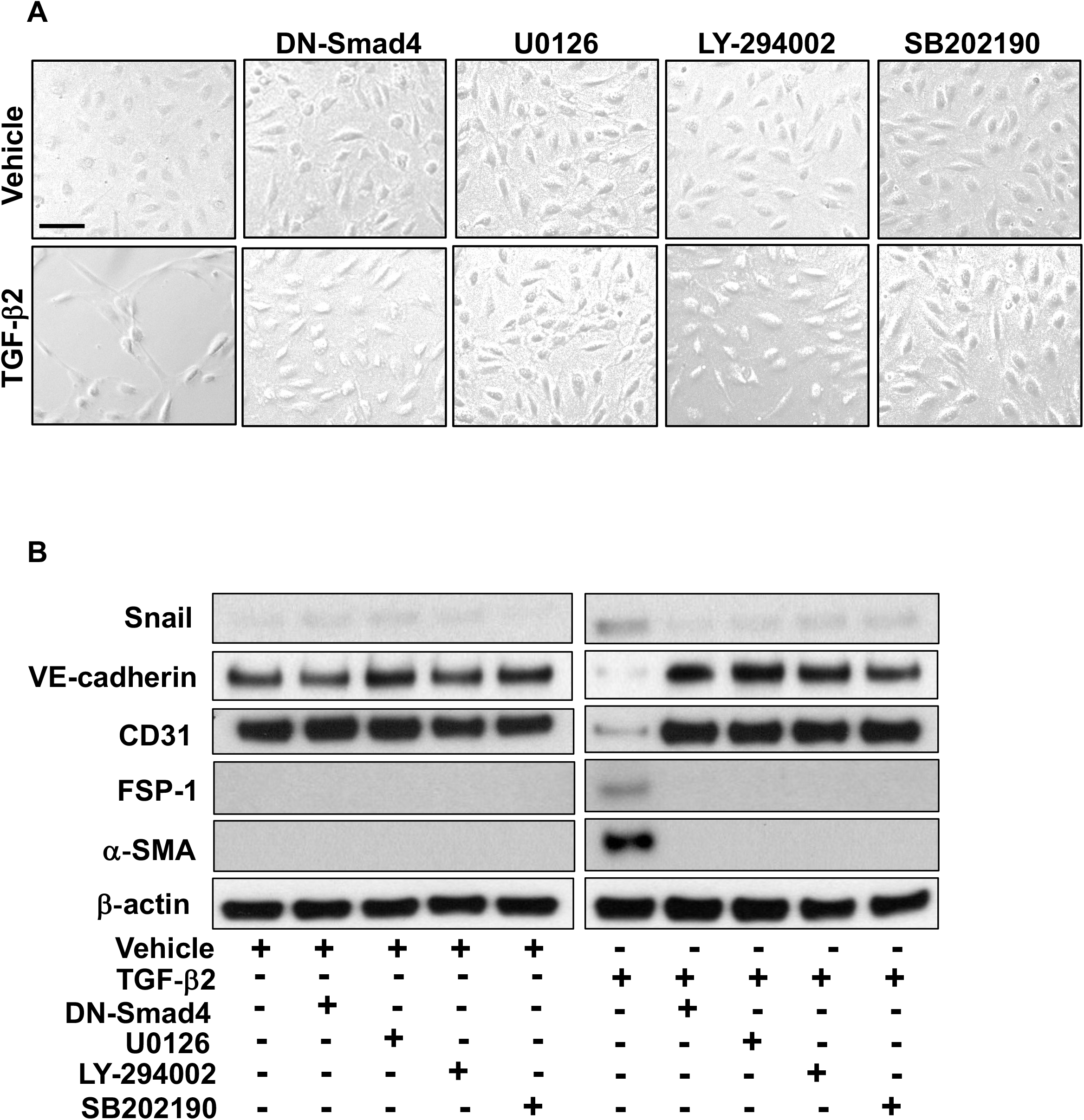
TGF-β2 promotes EndMT via Smad-dependent and Smad-independent signaling pathways. **A**. Representative bright field images of endothelial cells with the indicated treatment. Scale bar: 100μm. **B**. Immunoblotting for Snail, VE-cadherin, CD31, FSP-1, α-SMA, and β-actin (loading control). TGF-β2: 10ng/mL; Smad4 (DN-Smad4); MEK1/2 inhibitor (U0126), 10μM; PI3K inhibitor (LY294002), 50μM; p38 inhibitor (SB202190), 25μM.

**Figure 3.**
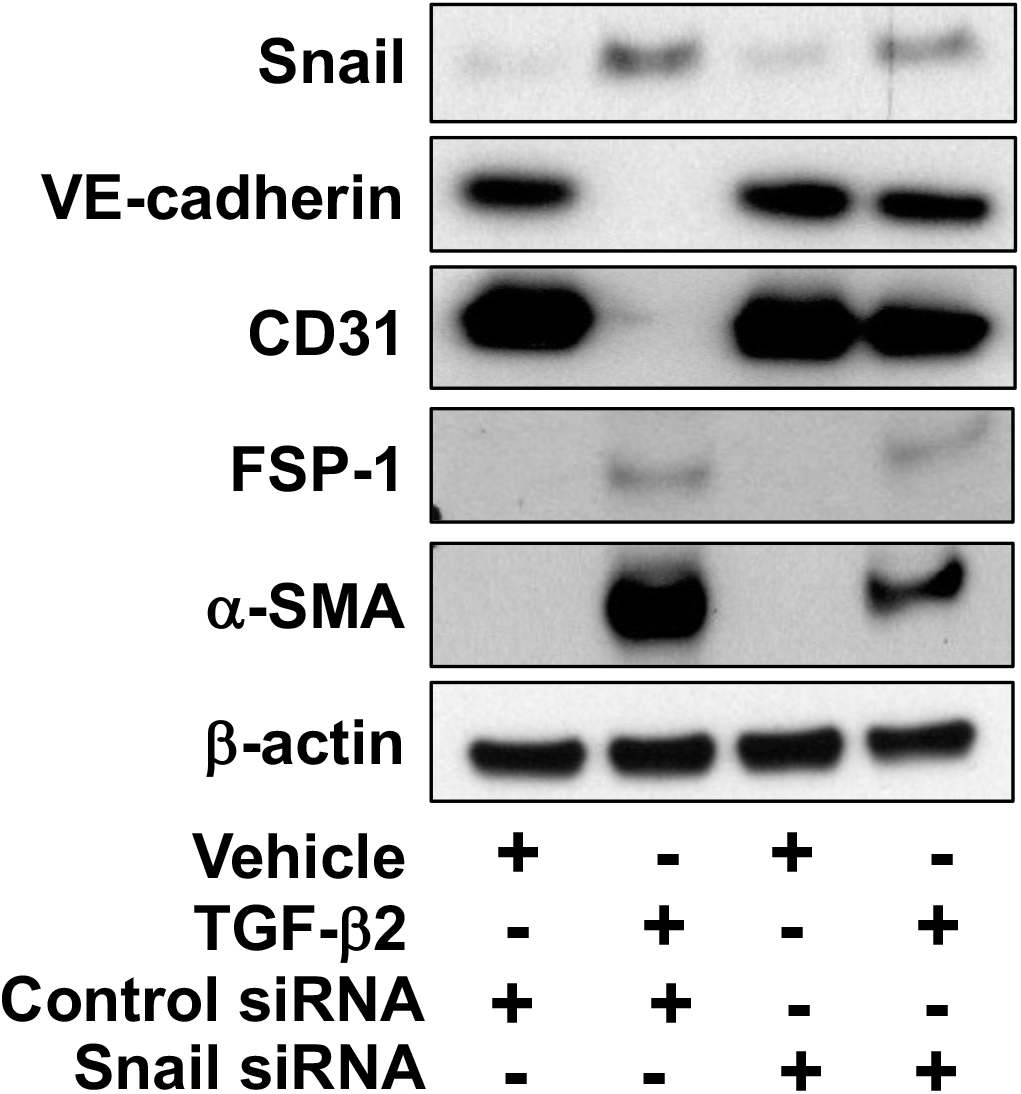
Snail activity is essential for TGF-β2-induced EndMT. Immunoblotting for Snail, VE-cadherin, CD31, FSP-1, α-SMA, and β-actin (loading control) in endothelial cells treated with or without TGF-β2 and with or without silencing of Snail using siRNA. TGF-β2: 10ng/mL.

**Figure 4.**
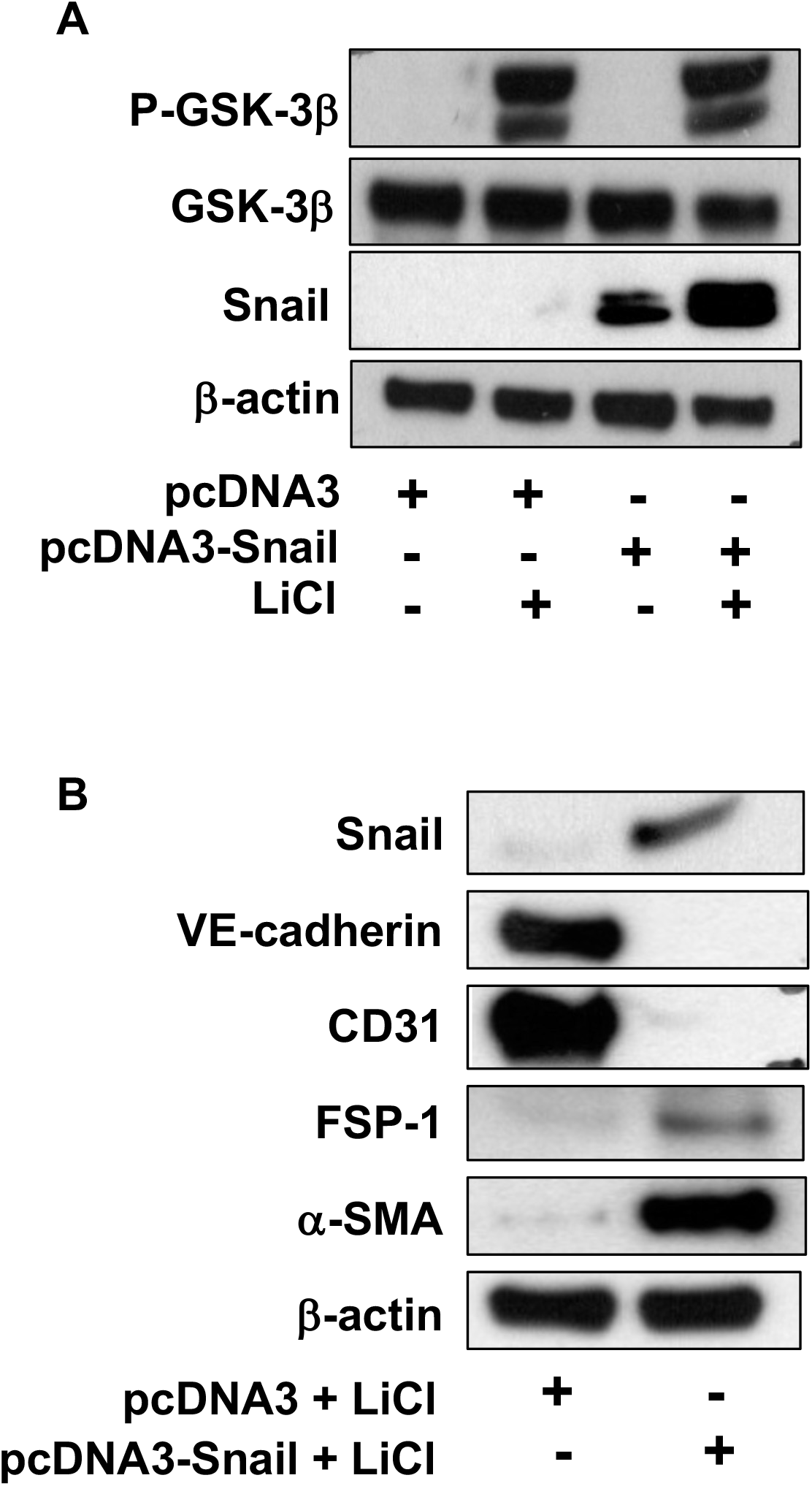
Snail overexpression is insufficient to induce EndMT. **A**. Immunoblotting for phosphorylated GSK-3β, total GSK-3β, Snail, and β-actin (loading control) in endothelial cells with Snail over expressing plasmic and with or without treatment for LiCl for GSK-3 inhibition. LiCl: 20mM. **B**. Immunoblotting for Snail, VE-cadherin, CD31, FSP-1, α-SMA, and β-actin (loading control) in endothelial cells treated with LiCl for GSK-3 inhibition and with or without overexpression of Snail. LiCl: 20mM.

## Discussion

The current study provides a comprehensive assessment of the reproducibility of one of the foundational mechanistic studies defining TGF-β2-induced endothelial-to-mesenchymal transition. Rather than evaluating reproducibility solely through direct experimental repetition, we combined two complementary approaches: systematic assessment of independent literature spanning fifteen years and contemporary laboratory replication of key biochemical findings. Our literature analysis demonstrates consistency in the principal mechanistic conclusions of our previously reported study^10^. Independent investigators reproduced activation of canonical Smad signaling together with ERK, PI3K/AKT, and p38 MAPK pathways downstream of TGF-β2, confirmed the central role of Snail as a regulator of EndMT, and further established the importance of GSK-3β-mediated regulation of Snail activity^11,12,18,22,29,34^. Importantly, these observations were reproduced in numerous endothelial cell populations originating from distinct vascular beds and across diverse pathological conditions, suggesting that the proposed signaling architecture represents a broadly conserved mechanism rather than one restricted to a particular experimental model.

The independent laboratory replication performed in this study provides an additional level of validation. Although experimental conditions inevitably evolve over time through changes in reagents, antibody lots, instrumentation, and laboratory personnel, the principal biochemical observations were reproducible using contemporary methodologies. This work also highlights an important distinction between technical reproducibility and conceptual reproducibility. Technical reproducibility addresses whether a specific experiment can be repeated under comparable conditions, whereas conceptual reproducibility examines whether the biological conclusions remain valid across independent investigators, experimental systems, and disease models. The latter may provide a particularly informative measure of the durability and generalizability of mechanistic discoveries in biomedical science.

Several limitations should be acknowledged. The literature analysis necessarily reflects the published scientific record and may therefore be influenced by publication bias. Furthermore, although multiple experiments were independently repeated, not every experiment from the original publication was reproduced experimentally. Nevertheless, the combination of direct laboratory replication and extensive independent validation across the literature provides compelling evidence supporting the robustness of the original mechanistic model. In conclusion, fifteen years of subsequent investigation demonstrate that the central signaling mechanisms governing TGF-β2-induced EndMT have proven highly reproducible across independent laboratories, experimental models, and disease contexts. Beyond validating a single study, this work illustrates how cumulative independent evidence can serve as an important and complementary measure of reproducibility for influential mechanistic discoveries in biomedical research.

## Acknowledgement

We acknowledge the use of ChatGPT (GPT-5.6 Sol) to assist with the writing and editing of the manuscript. Following use of this tool, the authors reviewed and edited the content as needed.

## Conflict of interest

The authors declare no competing interests.

## Supplementary Figure Legends

**Supplementary Figure 1**. Uncropped western blots for data shown in Fig. 1 (A) and Fig. 2 (B).

**Supplementary Figure 2**. Uncropped western blots for data shown in Fig. 3.

**Supplementary Figure 3**. Uncropped western blots for data shown in Fig. 4A (A) and Fig. 4B (B).

## Notes

### Competing Interest Statement

The authors have declared no competing interest.

