## Supplementary figures and images for "TGF-β2-induced Snail-mediated EndMT signaling relies on both Smad-dependent and independent pathways"

### Supplementary Figures

#### Supplementary Figure 1

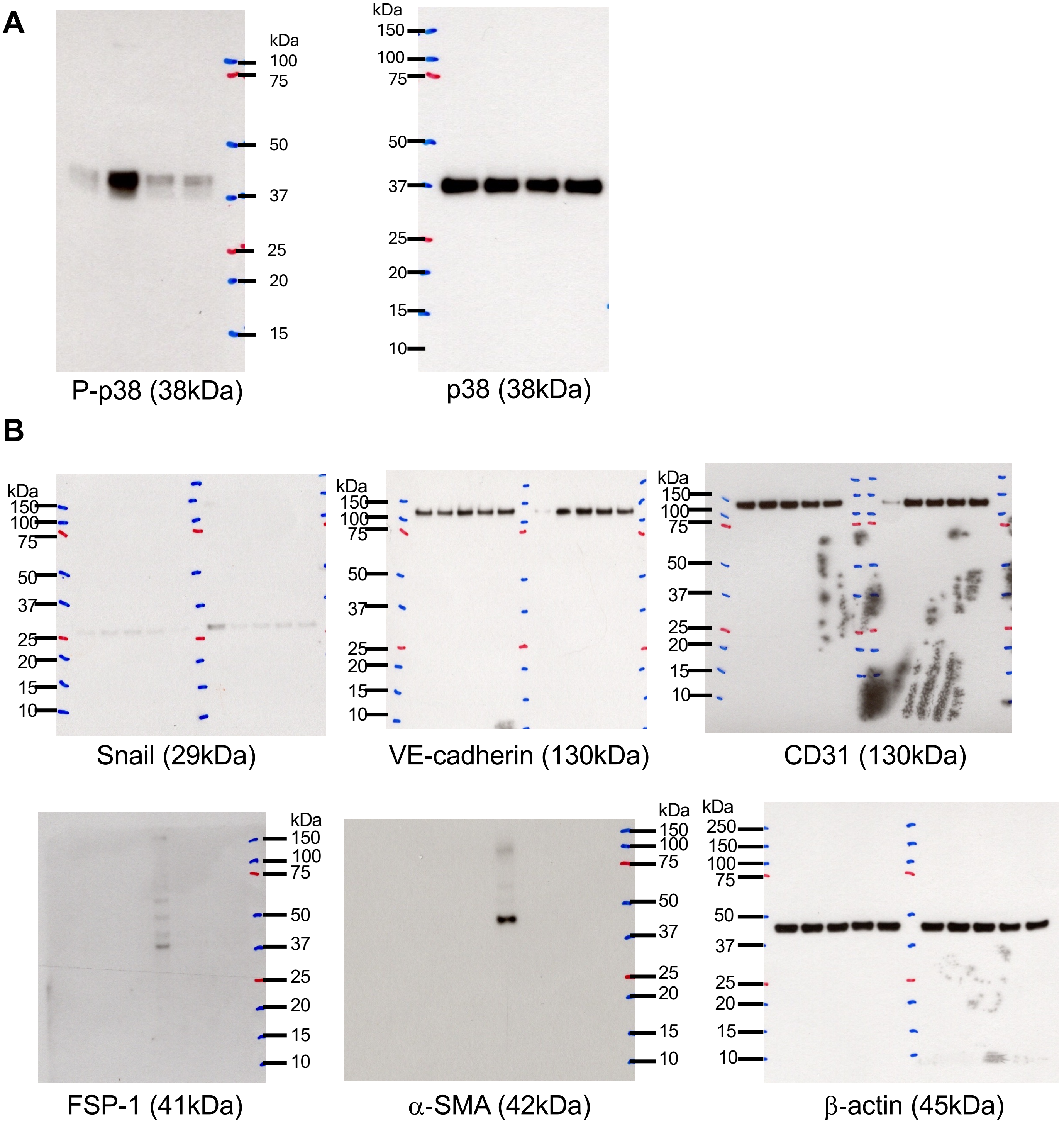

#### Supplementary Figure 2

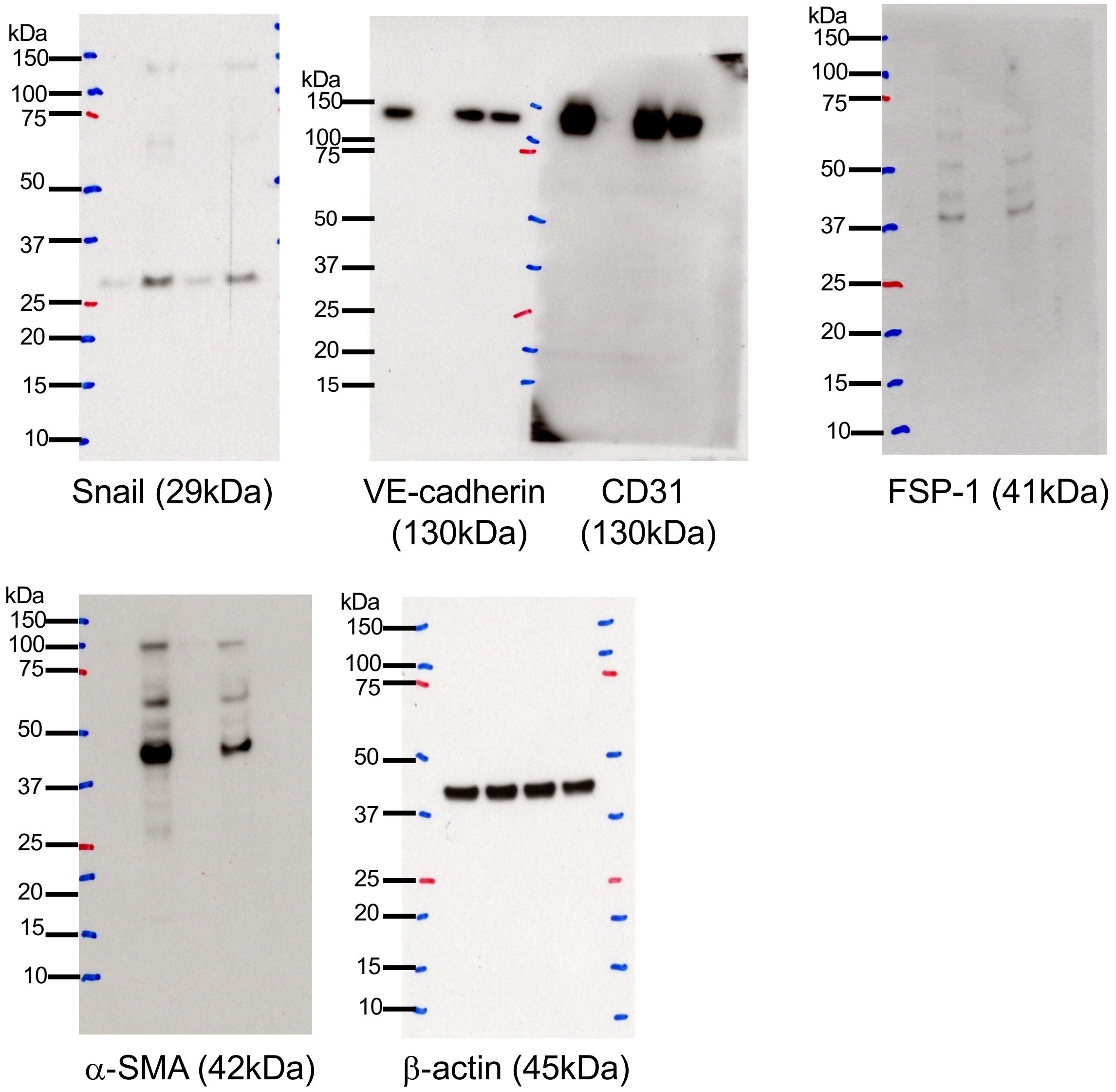

#### Supplementary Figure 3

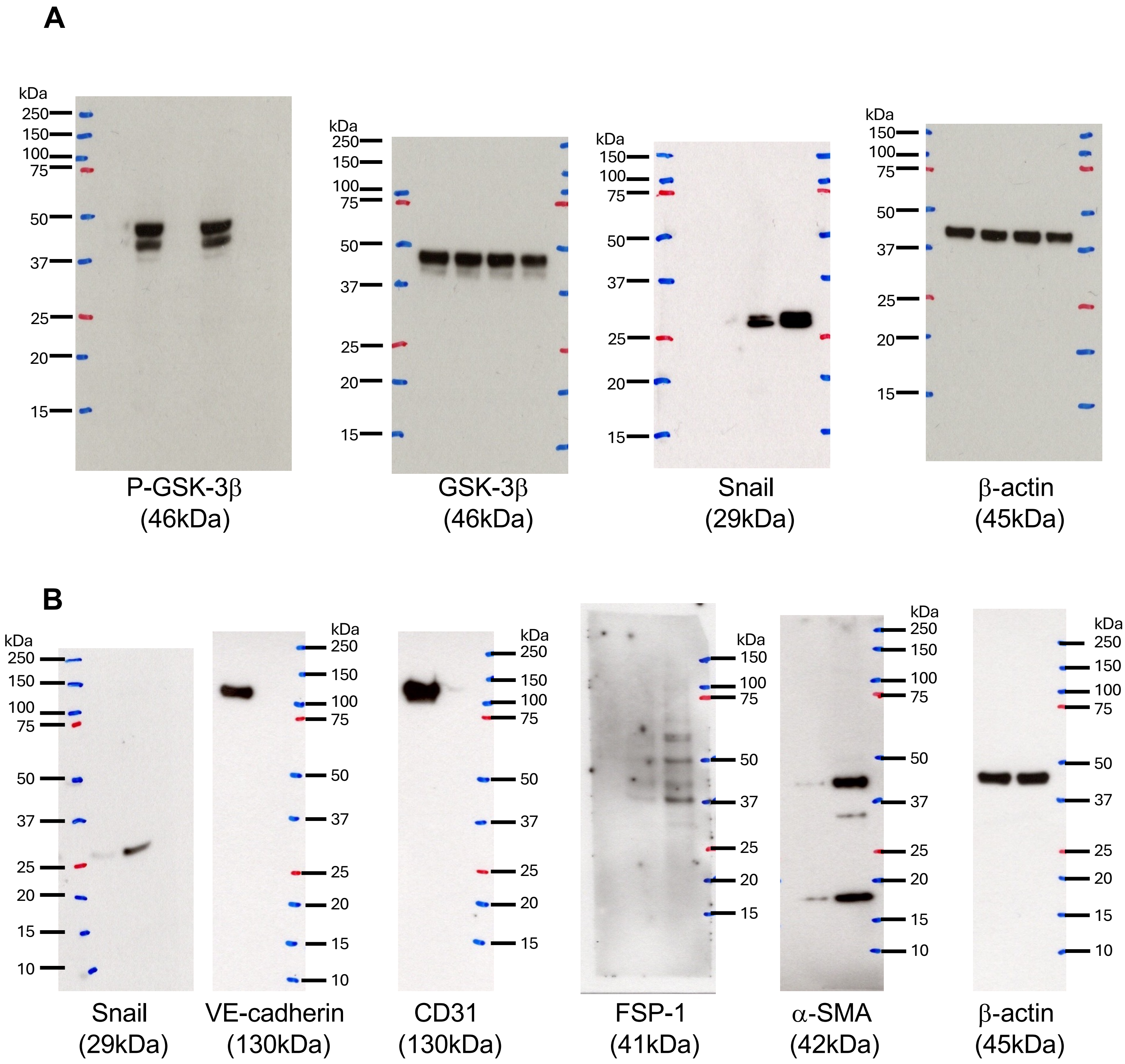
